# Heterogeneous and Surface-Catalyzed Amyloid Aggregation Monitored by Spatially Resolved Fluorescence and Single Molecule Microscopy

**DOI:** 10.1101/2022.10.05.510935

**Authors:** Xin Zhou, Anders Wilgaard Sinkjær, Min Zhang, Henrik Dahl Pinholt, Hanne Mørck Nielsen, Nikos S. Hatzakis, Marco van de Weert, Vito Foderà

## Abstract

Amyloid aggregation is associated with many diseases and may also occur in therapeutic protein formulations. Addition of co-solutes is a key strategy to modulate the stability of proteins in pharmaceutical formulations and select inhibitors for drug design in the context of diseases. However, the heterogeneous nature of this multi-component system in terms of structures and mechanisms poses a number of challenges for the analysis of the chemical reaction. Combining a spatially resolved fluorescence approach with single molecule microscopy and machine learning approaches, we disentangle the different contributions from multiple species within a single aggregation experiment. Moreover, we link the presence of interfaces to the degree of heterogeneity of the aggregation kinetics and retrieve the rate constants and underlying mechanisms for single aggregation events, providing a general tool for a comprehensive analysis of self-assembly reactions.

**Table of Contents:** 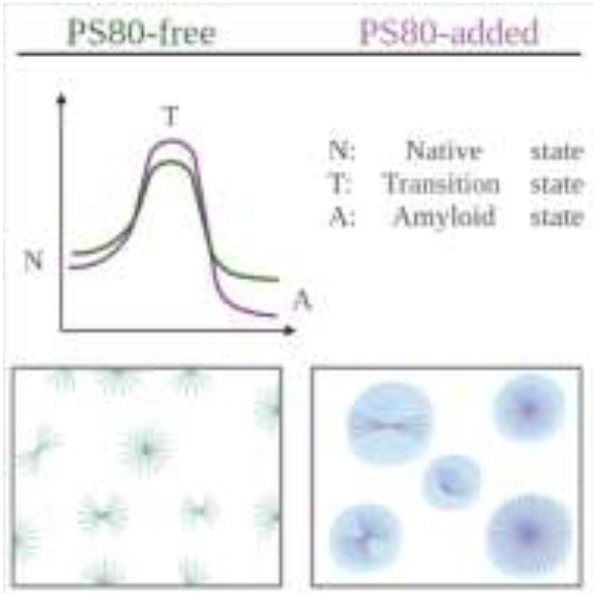

---

Deposition of amyloid protein aggregates in the form of fibrils is related to the onset of pathologies such as Alzheimer’s and Parkinson’s disease^[1-2]^, and diabetes Type II^[3]^. Moreover, protein aggregates can occur as an unwanted degradation product in protein and peptide pharmaceutical formulations^[4-5]^, and their occurrence can lead to lower efficacy of the therapeutic protein or peptide^[6]^ and increase the immunogenicity risk associated with use of the drug^[7]^.

Proteins may aggregate into different amyloid forms, spanning a broad range of characteristic sizes (from hundreds of nm to mm) and morphologies (from elongated linear structures and dense particles to core-shell architectures, named spherulites)^[8-10]^. Transition from the soluble state of the protein to the aggregated form is often accompanied by a transition from native-like structures to β-sheet-rich structures^[11-12]^. These multiple processes and morphologies are summarized in the generalized energy landscape for protein aggregation in which distinct energy minima represent the heterogeneous pool of aggregates that can be formed^[13]^.

One of the strategies to control the stability of a protein in solution — and in turn its aggregation propensity — is the addition of co-solutes^[14-16]^. For example, nonionic surfactants are often added in pharmaceutical formulations to prevent the therapeutic protein or peptide from aggregating^[17-18]^ or to minimize protein adsorption to interfaces^[18]^. Moreover, addition of surfactants, and in general co-solutes, alters the equilibria between the conformational states of the proteins and, consequently, modifies both protein-protein and protein-solvent interactions^[15-16]^. Addition of co-solutes may also introduce an enhanced variety of aggregation pathways, which eventually will result in the formation of a plethora of aggregate structures. Such heterogeneity makes the analysis of these systems very difficult when only using standard bulk assays, which will inevitably mask the broad spectrum of molecular mechanisms occurring at different time and length scales.

In a previous study, we identified a surprising effect of the commonly used surfactant polysorbate 80 (PS80) on the stability and aggregation propensity of insulin in solution^[19]^. Specifically, it was shown that PS80 slows down the thermal aggregation of insulin, but induced the formation of stable spherulites with an enhanced content of β-sheet structure and increased protein packing density^[19]^. This leads to our hypothesis that PS80 alters the mechanism of aggregate formation, the energetics related to the process, and the aggregate features, with these three aspects intrinsically related to each other.

In this work, we report a quantitative study on how the addition of PS80 modifies the energy landscape for the formation of insulin spherulites. Using a combination of calorimetry, fluorescence spectroscopy and electron microscopy, we estimate the energy barriers for aggregation of insulin in the presence of PS80 and relate these to the occurring morphologies. A method based on spatially resolved fluorescence spectroscopy is developed to evaluate the intrinsic heterogeneity of the aggregation process at the level of single aggregates. The latter allows us to identify multiple mechanisms during the reaction and avoid the loss of information, which occurs in bulk experiments. This is combined with our recently developed method based on real-time kinetic via photobleaching localization microscopy (REPLOM)^[20]^, by which the growth of spherulite over time can be extracted at the level of single molecules. We highlight an increase in heterogeneity of the spherulites formation kinetics, likely connected to a reduction of the initial surface-catalyzed aggregation in the presence of PS80.

With the aim of determining how PS80 affects the energy barrier associated with the transition of soluble insulin to the aggregated form, we used the amyloid-specific dye Thioflavin T (ThT)^[21]^ to detect the bulk aggregation kinetics at different PS80 concentrations and temperatures in the range 37–60 °C (Figure S1 in the Supplementary Information, SI). The acidic conditions used (see SI for details) promote the formation of insulin spherulites, as confirmed by the Maltese cross pattern^[22]^ obtained via cross-polarized light microscopy (Figure 1a and SI)

**Figure 1.**
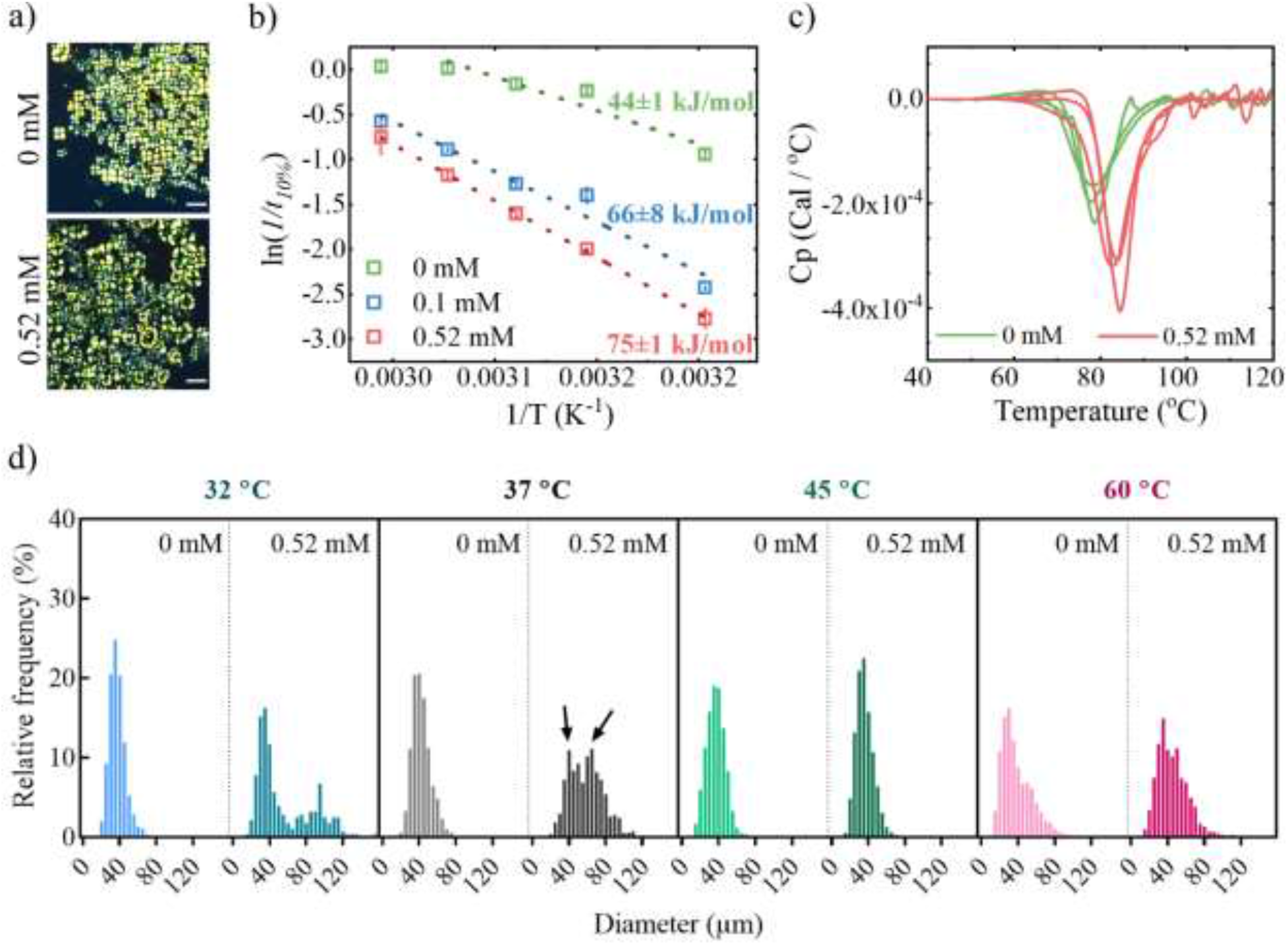
a) Polarized light microscopy images confirming the presence of insulin spherulites formed at 37 °C. 3 mg/mL insulin was dissolved in 20% (v/v) acetic acid with 0.5 M NaCl, in the absence (top) or presence (bottom) of 0.52 mM polysorbate 80 (PS80). b) Arrhenius plots obtained from Thioflavin T (ThT) kinetics (see SI and Figure S1), showing ln 1/*t*_10%_ as a function of reciprocal of the temperature (37 °C–60 °C) and PS80 (0–0.52 mM). *t*_10%_ is the time required to reach 10% of the kinetics completion (mean±SD, *n*=4, except 0 mM at 50 °C, and 0.1 mM at 37 °C where *n=*3; *n* represent the number of replicates). The dotted lines are the linear fits to the data from which energy barriers were calculated. c) Differential scanning calorimetry scans of 3 mg/mL insulin in 0 mM and 0.52 mM PS80 solutions in the above-mentioned buffer. 3 replicates per sample are included in the figure. d) The diameter distribution of spherulites formed at 32 °C, 37 °C, 45 °C, and 60 °C with different amounts of PS80 added. The arrows show the two distinct peaks in the distribution.

The kinetics show the well-known sigmoidal profile (Figure S1 in SI) and allow estimation of the time at which the amyloid aggregation starts. 1/*t*_10%_, the reciprocal of the time at which 10% of the aggregation reaction is complete ^[23-24]^ is defined as a model-free estimate of the rate constant for the nucleation process. When reported in an Arrhenius plot (1/*t*_10%_ vs. 1/T), the data obtained at 0.1 and 0.52 mM PS80 in acetic acid solutions show linear trends in the entire temperature range (blue and red symbols, respectively, in Figure 1b). In absence of PS80, linearity only holds in the range 37-55 °C and not at the highest temperatures (green symbols, Figure 1b). Interestingly, this behavior may be due to a subtle balance between association and dissociation processes taking place simultaneously^[25]^. At high temperatures, the rate of dissociation may be comparable to the association rate, resulting in the absence of a temperature dependency for the overall rate of the process expressed by 1/*t*_10%_. This effect is not detected in the presence of PS80, in agreement with our previous observation that PS80 induces a higher stability of the aggregates^[19]^, likely minimizing the impact of the dissociation reaction.

Evaluation of the bulk activation energies of the amyloid reaction indicates a clear effect of the PS80. In the absence of PS80, an activation energy of 44±1 kJ/mol was observed, while values of 66±8 kJ/mol and 75±1 kJ/mol were found for 0.1 mM and 0.52 mM PS80, respectively (Figure 1b). Differential scanning calorimetry data in the range 10-120 °C show that, both in absence and in presence of 0.52 mM PS80 in the acetic acid solution, an exothermic transition attributed to the aggregation of insulin^[26]^ dominates the thermograms. The addition of PS80 induced a shift in the position of the exothermic peak from ∼78 °C to ∼84 °C (Figure 1c), confirming an enhanced energy barrier for aggregation in the presence of PS80. PS80 was previously reported to inhibit the shaking-induced aggregation of recombinant human IL-2 mutein^[27]^. Indeed, polysorbates are known to occupy the air-liquid or solid-liquid interfaces, reducing surface-catalyzed protein unfolding and aggregation and stabilizing the protein in solution^[17-18, 28]^. We may infer a similar effect from the observations in our system^[19]^, leading to an increase in the aggregation energy barrier in the presence of PS80 (Figure 1b). Moreover, our previous study suggested that PS80 molecules may also interact with insulin hydrophobic regions, potentially reducing the chance of inter-molecular hydrophobic interactions and aggregation^[19]^. This aspect may reduce the number of molecules available for aggregate formation and reduce the number of effective attempts to overcome the energy barrier for the process^[29]^.

The size distribution of spherulites was also estimated together with their abundance from image analysis of optical microscopy data (Figure S2-S5, and details in SI and in reference^[29]^). This approach may give information on the efficiency of the nucleation process^[29]^. In the range of 45-60 °C, the spherulite sizes and numbers did not significantly vary upon addition of PS80 (Figure 1d and Table 1). In contrast, at 32 °C and 37 °C, the presence of PS80 lead to the appearance of a group of spherulites with larger sizes, showing a bimodal distribution (Figure 1d) and a decrease in the number of detected spherulites (Table 1). Thus, in the absence and presence of PS80, the median of the monomodal size distributions at 32 °C and 35 °C changed from 35.6 μm and 41.0 μm to bimodal distributions with medians of 35.0 μm and 93.3 μm, and 42.1 μm and 68.6 μm, respectively (Table 1). A similar trend was previously observed as a function of salt concentration^[29]^. In our specific case, the soluble-to-aggregate conversion rate is independent of the PS80 concentration and close to 100% in all cases (Table S2 in SI). These observations indicate that the enhanced barrier induced by PS80, combined with the low temperature of incubation, reduces the number of nucleation events and, ultimately, the number of spherulites. This means that more soluble protein molecules are available for the actual growth of the spherulites in the presence of PS80 compared to the conditions in the system in the absence of PS80 and/or when incubated at high temperature. This results in fewer spherulites reaching a larger size in the presence of 0.52 mM PS80 at both 32 °C and 37 °C

**Table 1.** The count number of spherulites and the median (μm) of the diameter distribution of counted spherulites summarized in Figure 1d. 3 mg/mL insulin was dissolved in 20% (v/v) acetic acid and 0.5 M NaCl, with addition of no or 0.52 mM polysorbate 80 (PS80) at different temperatures.

| Temperature | 32 °C |  |  | 37 °C |  |  |
| --- | --- | --- | --- | --- | --- | --- |
| [PS80] | 0 mM | 0.52 mM |  | 0 mM | 0.52 mM |  |
| Number | 1418 | 283 |  | 1017 | 541 |  |
| Median, $\mu\text{m}$ | 35.9 | 35.0 | 93.3 | 41.0 | 42.1 | 68.6 |
| Temperature | 45 °C |  |  | 60 °C |  |  |
| [PS80] | 0 mM | 0.52 mM |  | 0 mM | 0.52 mM |  |
| Number | 1438 | 1998 |  | 986 | 1124 |  |
| Median, $\mu\text{m}$ | 36.6 | 34.8 | | 34.6 | 42.7 | |

Taken together, the data in Figure 1 highlight a scenario in which PS80 increases the energy barrier for insulin aggregation, particularly affecting the abundance and size of the formed spherulites at the two lowest temperatures investigated.

Energy barriers of the amyloid aggregation kinetics, and in turn the rate constants of the process, are dramatically dependent on initial protein concentrations, homogeneity of the chemical species in solution, geometry of the sample holder, and protein interactions with both solid and liquid interfaces. The above-mentioned factors are also recognized as potential sources of the intrinsic variability of the kinetic profiles. Specifically, Grigolato and Arosio recently showed that — given identical conditions in connection to the interfaces of the sample holder and absence of pre-formed seeds^[30]^ — the variability of the aggregation profiles is particularly sensitive to the initial monomer concentration^[31]^. The non-ionic surfactant PS80 was previously indicated to prevent protein adsorption to container surfaces and the air–water interface through preferential adsorption^[32-33]^. Moreover, insulin aggregation is reported to be initiated via surface-catalyzed mechanisms^[34]^. Based on these reports, we hypothesize that, in the presence of PS80, the surface-catalyzed effect may be strongly reduced, favoring an aggregation process mainly dominated by bulk aggregation. Bulk aggregation can be highly heterogeneous in space and time^[35-36]^, which is reflected in a pronounced variability of the kinetic profiles even within a single sample. To verify this, we designed a spatially resolved ThT fluorescence assay during the incubation of insulin in aggregation conditions. Briefly, within a single well in a 96-well plate, 21 different areas were scanned and 21 ThT temporal profiles were extracted (Figure 2a and b). In the absence of PS80, the ThT profiles overlap, showing a negligible variability in terms of nucleation time (*t*_10%_) between the traces (Figure 2b). At increasing concentrations of PS80, the ThT trace variability consistently increased, showing a pronounced difference between different areas of the well at the highest PS80 concentration (Figure 2b).

**Figure 2.**
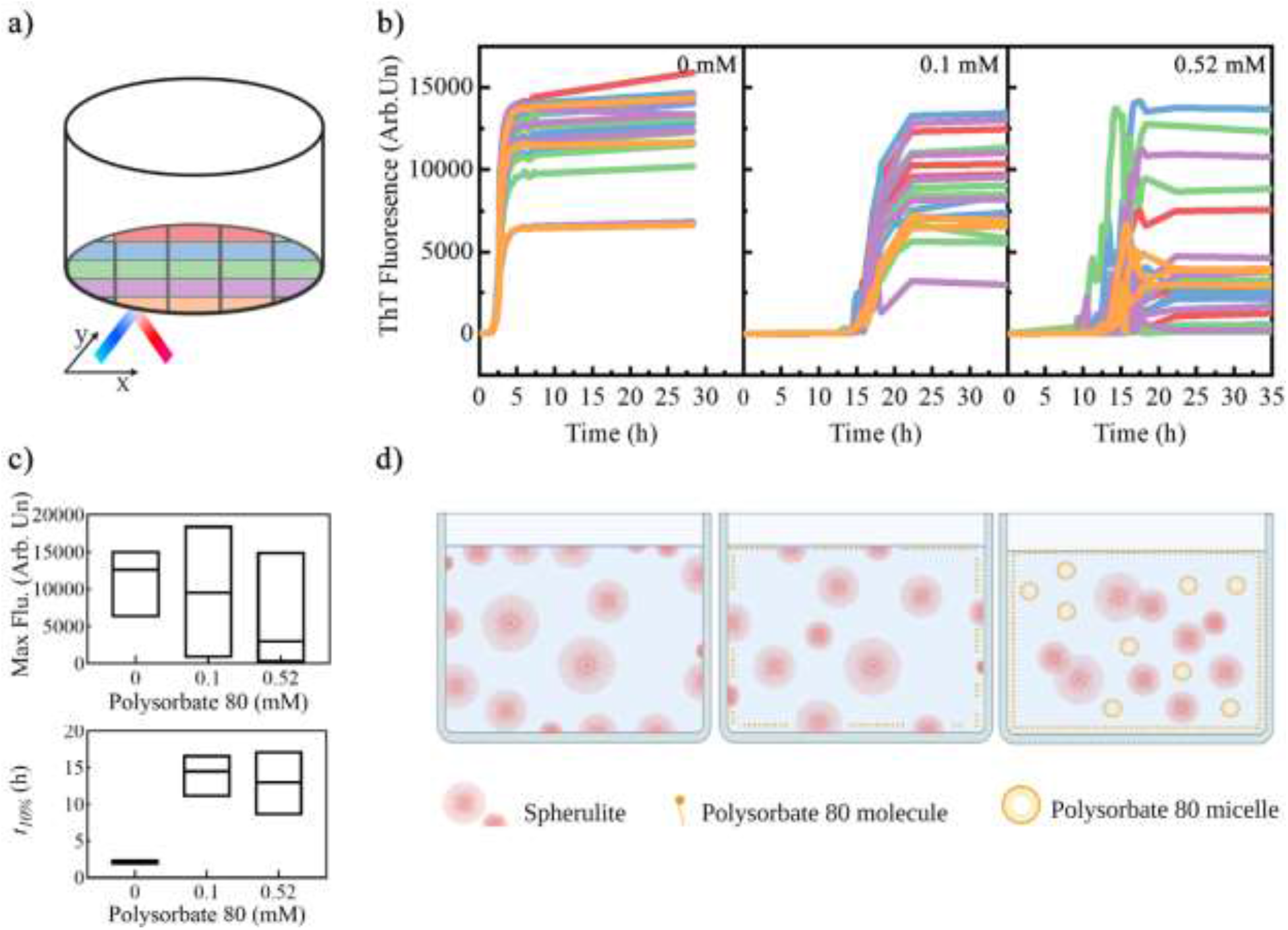
a) Illustration of grid localization in one well for the kinetic studies. b) Example of grid kinetics of insulin amyloid formation at 0 mM, 0.1 mM and 0.52 mM of polysorbate 80 (PS80). Traces are color-coded according to the illustration in a) and represent kinetic profiles being detected at different localization in a well. c) Maximum fluorescence values (MFV) and the time needed for reaching 10% of MFV (*t*_10%_). The lower and upper limits of the boxes represent minimum and maximum values observed, respectively. The line in the boxes represent the median of the distribution. The analysis was performed on grid kinetics from 4 wells. The distributions of the MFV and *t*_10%_ were calculated from more than 72 kinetic traces. d) Illustration of the effect of PS80 molecules on insulin amyloid spherulites formation, created with BioRender.com.

In the presence of 0.52 mM PS80, no ThT signal was observed in certain areas even on a time scale of 10 hrs (Figure 2b). This increased variability is reflected in the box plots of the distribution of the maximum fluorescence value (MFV) and *t*_10%_ of the kinetics for each PS80 concentration. Both the MFV and the *t*_10%_ are distributed more broadly when the aggregation reaction occurred in the presence of PS80 (Figure 2c). We attribute this variability to the known heterogeneity in space and time of the bulk nucleation and growth process^[35-36]^. Indeed, at the high surfactant concentration, PS80 may occupy both air-liquid and liquid-solid interfaces in the sample holder, strongly reducing surface-catalyzed aggregation in favor of bulk aggregation (Figure 2d). In contrast, in the absence of PS80, the catalytic effect of the surface was prominent, inducing a fast aggregation within the entire sample volume, as can be observed by completion of the reaction in less than 2 hrs (Figure 2b). Surface effects at the level of biological membranes in vivo or the vessel and the air/water interface in vitro are indeed considered as modulators of the aggregation reproducibility^[37]^. These observations are also in line with the fact that the variability of the macroscopic kinetic traces increases linearly with the duration of the aggregation process^[31]^. In our case, the balance between bulk and surface-catalyzed aggregation determines the duration of the process and, in turn, the reproducibility of the process.

With the aim of detecting how PS80 affects the morphological evolution of spherulites and their mechanism of formation, we first performed a time-lapse analysis using transmission electron microscopy (TEM, Figure S6 in SI). To avoid potential artefacts due to the long duration of the aggregation kinetics at the highest PS80 concentration and to have a suitable timescale of the aggregation for imaging analysis, we limited the study to 0 and 0.1 mM PS80 and incubation at 45 °C. By TEM, we detected a delay in the appearance of amyloid-like structures in samples when PS80 was added. However, it is still unclear how individual spherulites are formed and affected by PS80.

The nucleation theory has been accepted as a description of the three-step kinetic profile of the amyloid fibril formation^[2, 38-39]^ with the nuclei formation mainly occurring during the lag phase^[40]^. Subsequently, protein monomer additions to the nucleus and secondary nucleation processes determine the growth of the fibrils^[39]^. For spherulites, a compact core via a multifractal pattern was originally suggested^[41]^, in which branched protein material isotropically grows on the core. Recently, we have developed an approach for REal-time kinetics via Photobleaching LOcalization Microscopy (REPLOM)^[20]^. This allows one to track the aggregation kinetic traces while simultaneously recording the morphological development of spherulites in real time and at single aggregate level (see videos and details in SI and reference^[20]^). Using this approach, we reported anisotropic spherulite growth patterns, highlighting a high heterogeneity in the growth mechanisms^[20]^. More specifically, we reported an initial preferential direction of the growth from the core, followed by branching leading to the formation of mature spherulites^[20]^.

We employed REPLOM to study the specific effect of PS80 on the spherulite growth at single aggregate level, in absence of PS80 and at 0.1 mM PS80 (see SI for details). For both samples, we identified groups of spherulites formed through either isotropic or the anisotropic growth from the core of the spherulites (see videos in SI and Figure 3a-b for snapshots). Using homemade software based on Euclidian minimum spanning tree and machine learning clustering^[42-44]^, we calculated the growth rate of each spherulite morphology. In the first case, a single rate constant k was extracted by linear fitting of the initial kinetics traces followed by a plateau up to approximately 200 frames (100 min) (Figure 3a and details in SI)^[20]^. The anisotropic growth kinetics displayed an initial growth phase with a rate constant k_1_ associated with a linear growth of the aggregate, whereas a second phase with a higher rate constant k_2_ was related to the branching of the protein material (Figure 3b).

**Figure 3.**
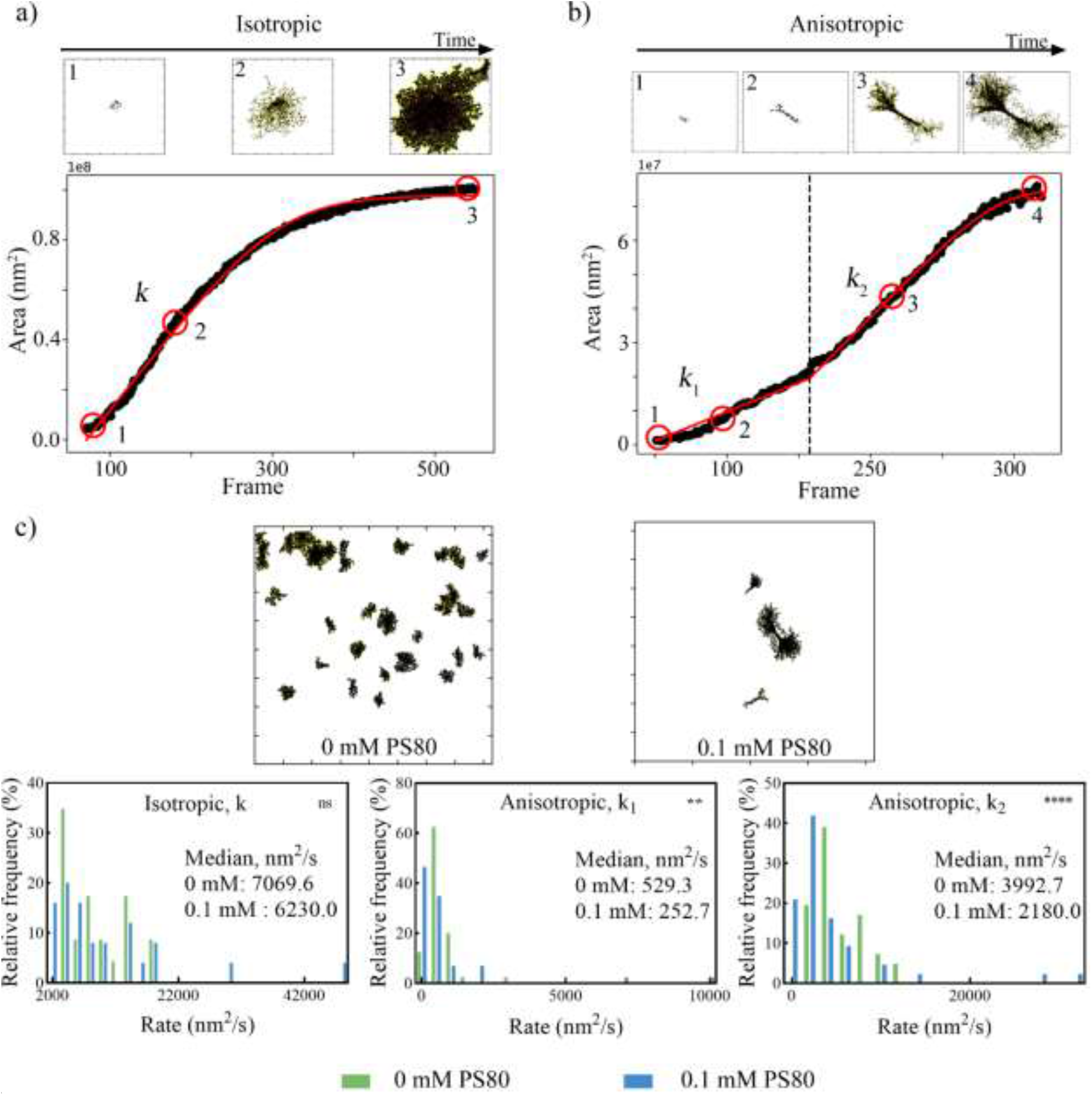
Spherulites growth kinetics recorded in real time via photobleaching localization microscopy (REPLOM). (a) isotropic growth and (b) anisotropic growth. Linear fits with a sigmoidal saturation to individual spherulites growth are shown in (a) and (b) by red lines. The numbers marked on the growth curve in (a) and (b) refer to the growth stages of the spherulite at the frames shown above the panels. (c) Top: Overview images (81.92 μm×81.92 μm) of the aggregates detected in a single area after the REPLOM experiment, without and with 0.1 mM of polysorbate 80 (PS80). Bottom: Rate constants distribution according to the fitting obtained from REPLOM. Non-significant differences (ns, p>0.05) and significant differences (**, p=0.0024 and ****, p<0.0001) are indicated (t-tests). 23 (0 mM PS80) and 25 (0.1 mM PS80) spherulites were included in the analysis for the isotropically grown spherulites, while 41 (0 mM PS80) and 43 (0.1 mM PS80) for the anisotropically grown ones. Experiments were conducted by dissolving 3 mg/mL insulin in 20% (v/v) acetic acid and 0.5 M NaCl at 45 °C, with 0.1 mM or without the addition of PS80.

The distribution of growth rates for different spherulites is presented in Figure 3c (bottom). The distribution of rates is highly polydisperse as expected. Interestingly, PS80 only slightly affected the rate of isotropic growth k, reducing the rate by ∼11.9%. For the spherulites growing anisotropically, the presence of the surfactant reduced k_1_ and k_2_ in a statistically significant manner (∼52.3% and ∼45.4%, respectively). Almost all available insulin assembled into aggregates in both samples (Table S2), independent of the presence of PS80. Moreover, fewer aggregates per unit area were detected in the presence of PS80 (representative micrograph in Figure 3c, top).

At present, we cannot explain the physical laws governing the origin of the observed isotropic and anisotropic growth. However, our experimental results unambiguously show the existence of a heterogeneous population of aggregate cores in terms of their available sites for aggregation on their surface. Specifically, for the isotropic growth one can infer that the entire surface of the core is available for binding to protein molecules (Scheme 1a, top) generating the growth pattern observed in Figure 3a. In contrast, apparently only a limited part of the core surface (Scheme 1b, top) is active/available when the spherulites grow anisotropically (Figure 3b). We postulate that in the presence of PS80, the free surfactant, i.e. PS80, not covering liquid-solid and liquid-air interfaces, may occupy some of these available binding sites, preventing the insulin assembly and delaying the aggregation process (Scheme 1a and 1b, bottom). In principle, this effect should take place for both isotropic and anisotropic growth pathways, but the impact will depend on the ratio between number of available binding sites and amount of PS80 molecules, as well as a potential different hydrophobicity or structure of these binding sites. Indeed, when the isotropic growth is dominant, we observe only a slight decrease in the rate constant k when PS80 is present (∼11.9%), while a significant decrease is observed for k_1_ and k_2_ during anisotropic growth (∼52.3% and ∼45.4, respectively). The difference in available sites for the growth in the two scenarios may explain these observations. In the case of anisotropic growth, the used PS80 concentration likely guarantees a higher PS80 occupancy of the available sites compared to the isotropic case, drastically reducing the rate of the process. This would explain the data in Figure 3c (bottom), and the more pronounced effect of PS80 on the anisotropic growth.

**Scheme 1.**
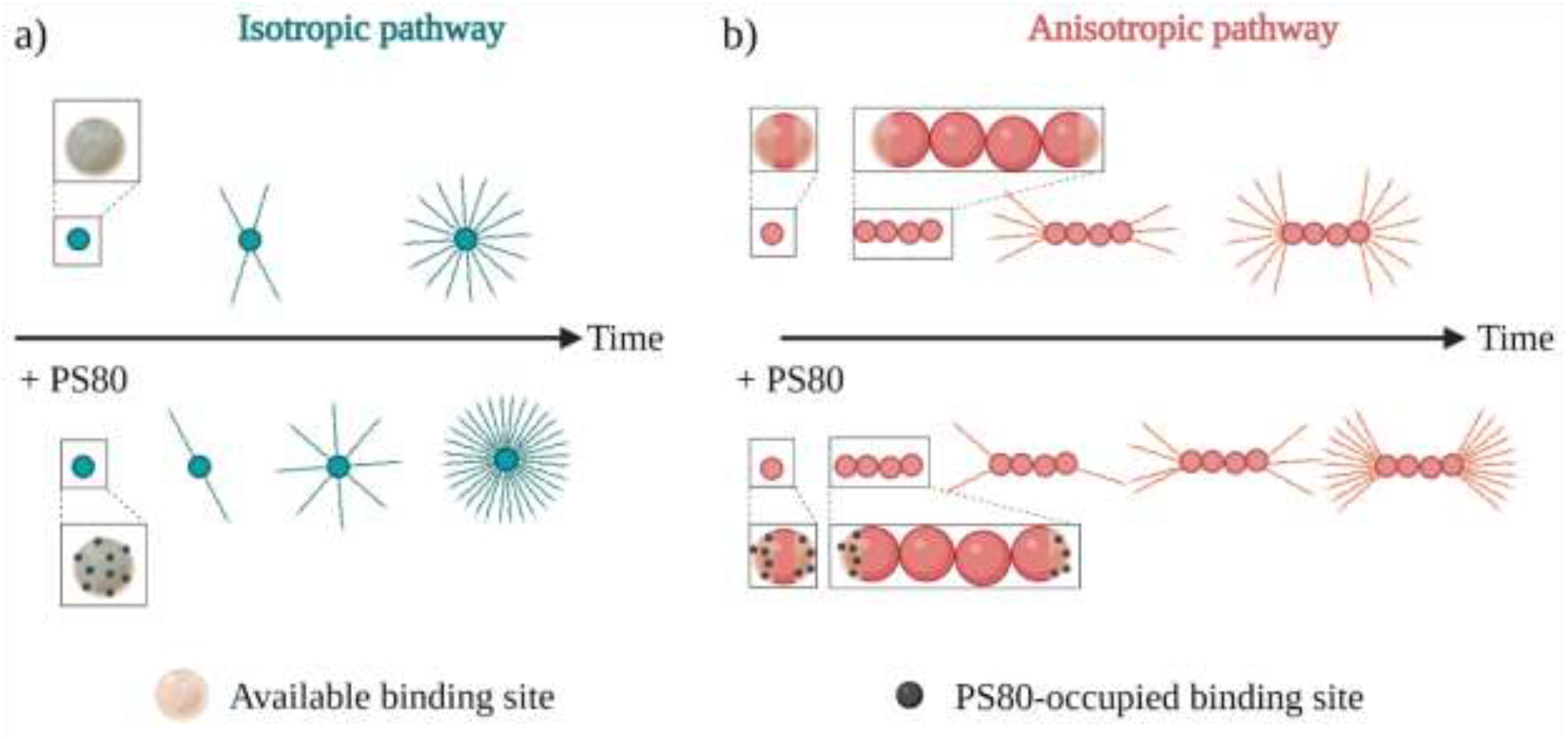
Illustration of spherulites formation. (a) The isotropic pathways of a spherulite formation in the absence of PS80 (top) and the presence of 0.1 mM PS80 (bottom) are shown. The isotropic pathway starts with the formation of a core from which fibril-like structures expand as insulin molecules adds to the binding sites in all directions. PS80 addition blocks some of the binding sites (bottom) thereby delaying the growth. (b) The anisotropic growth consists of two phases: The first phase is identified as the core formation and elongation along a preferential axis. Following nucleation, the elongated structures spread from the binding sites at the ends of the axis during the second phase. PS80 acts by occupying some available binding sites (bottom). The ratio between PS80-occupied binding sites and total available sites is higher in the anisotropic growth pathway compared to the isotropic growth pathway. This leads to a more pronounced decrease of the rate for the anisotropic growth of spherulites. The figure was created using BioRender.com.

In conclusion, we show that PS80 increases the energy barrier related to the conversion of soluble human insulin into amyloid-like spherulites. Using a spatially resolved fluorescence approach, we attribute the increase of the barrier in presence of PS80 to a shift from surface-catalyzed aggregation to bulk aggregation, which is reflected in an enhanced variability of the kinetic traces within a single sample volume. Moreover, freely available PS80 molecules in solution alter the rate constants for the spherulite growth at the single molecule level. We disentangled the role of PS80 to the growth of two types of spherulites: Isotropically- and anisotropically-grown spherulites, highlighting the need to consider the ratio between the number of PS80 molecules and the available sites for aggregation on the initially formed cores for correctly extracting the aggregation mechanisms. The synergy between suppression of the surface-catalyzed aggregation by PS80 and an effect of freely available PS80 molecules in solution is responsible for the overall increase of the energy barrier of insulin towards aggregation, providing a connection between microscopic mechanisms and thermodynamics of the chemical reaction.

Our findings are pivotal for studies on the effect of co-solutes on the aggregation of both biopharmaceuticals and medically-relevant proteins. Identifying the contributions of different aggregates within a single reaction may indeed provide a more comprehensive understanding of the mechanisms involved and result in a more efficient design and development of active molecules for stable drug formulations and medical treatments. Finally, our work promotes the use of spatially resolved ThT fluorescence for a rapid and high throughput evaluation of the kinetic variability of protein aggregation reactions, giving insights on the mechanisms ruling the processes.

## AUTHOR INFORMATION

VF conceived the idea, planned and managed the research activities. XZ, MZ and AWS did the experimental work. XZ, AWS, and HDP analyzed the data. XZ, MZ, NSH, MvdW, and VF took part in the data interpretation. XZ, MvdW and VF drafted the paper. AWS, MZ, HDP, HMN and NSH revised the manuscript.

The authors declare no competing financial interests.

## ACKNOWLEDGMENT

V.F. and X.Z. acknowledge China Scholarship Council (Project Number: 201709110108) for funding the project. V.F. and X.Z. acknowledge Villum Fonden for supporting the project via the Villum Young Investigator grant “Protein Superstructures as Smart Biomaterials (ProSmart)” 2018−2023 (Project Number: 19175). The authors acknowledge the Core Facility for Integrated Microscopy, Faculty of Health and Medical Sciences, University of Copenhagen for access to the transmission electron microscope. The authors kindly thank the Carlsberg Foundation Distinguished Associate Professor Program (CF16-0797) and (CF21-0499) to NSH, the Lundbeck foundation grant (R250-2017-1293 and R346-2020-1759) and the Carlsberg foundation grant (CF21-0659) to MZ. The authors kindly thank the Danish Research Council for Technology and Production Sciences for funding Centrifuge 5417R, Villum Fonden (19175) that funded the CLARIOstar plate reader and Shimadzu UV meter and Villum Fonden (19175), Novo Nordisk Foundation (NNF16OC0021948) and Lundbeck Foundation (R155-2013-14113) for funding the Leica DMi8 microscope. The authors thanks Dr. Jijo J Vallooran (University of Copenhagen) for the support on the transmission electron microscopy experiments.

