## Supplementary information for "Heterogeneous and Surface-Catalyzed Amyloid Aggregation Monitored by Spatially Resolved Fluorescence and Single Molecule Microscopy"

### Materials and Methods

#### Materials

Human insulin (≥95%), Thioflavin T (ThT, ≥ 65%), and polysorbate 80 (PS80) were obtained from Sigma Aldrich (Saint Louis, MO, USA). Acetic acid (≥ 98%) was obtained from VWR Chemicals (Leuven, Belgium). Sodium hydrochloride (99.0%) was obtained from Th. Geyer, CHEMSOLUTE (Roskilde, Denmark).

#### Thioflavin T Fluorescence (ThT) Kinetics

3 mg/mL (0.52 mM) insulin was dissolved in 20% (v/v) acetic acid and 0.5 M NaCl with or without PS80. The concentration of PS80 was 0 mM, 0.1 mM, or 0.52 mM. The pH was ~1.7 for all test samples and no further pH adjustments were done. Samples were filtered before inducing aggregation through a 0.22 μm cellulose acetate (CA) filter (Labsolute, Th.Geyer, Renningen, Germany). The insulin concentration estimated via NanoDrop 2000C (Thermo Scientific, Wilmington, NC, USA) by using 1.0 $(mg/mL)^{-1}cm^{-1}$ as the extinction coefficient at 276 nm. The insulin concentration can be found in Table S1. 1 mM of ThT stock solution was prepared by dissolving ThT powder in purified water and filtering through a 0.22 μm CA filter. The concentration was determined using the Nanodrop 2000C at 412 nm by using 36,000 $M^{-1}$ $cm^{-1}$ as the extinction coefficient[1]. The final concentrations of ThT in the protein solutions was 20 μM. ThT fluorescence was recorded during the aggregation reaction as a function of time on a Clariostar plate reader (BMG Labtech, Ortenberg, Germany) in 96-well polystyrene plates (Nunc, Thermo Fisher, Rochester, NY, USA). Each well contained 200 μL solution. A polyolefin film (Thermo Fisher, Rochester, NY, USA) was used to prevent evaporation during the measurements. Samples were incubated at 37, 45, 50, 55, and 60 °C without shaking until a plateau of the

fluorescence signal was reached for all samples. The emission intensity at 480 nm was recorded upon excitation at 450 nm every 309 s throughout the incubation time. Kinetics are shown in Figure S1. The Arrhenius plot was presented by plotting ln($1/t_{10\%}$) versus the reciprocal of the temperature, where $t_{10\%}$ represents the time needed to reach 10% of the maximum fluorescence value.

**Table S1**. Insulin concentration after filtration at different concentrations of polysorbate 80 (PS80) and prior aggregation reaction (Mean ± SD, n=3, N=2, *n* stands for different samples measured in one batch, while *N* indicated the number of the different batches).

| PS80, mM | Insulin, mg/mL |
|---|---|
| 0 | 2.95 ± 0.05 |
| 0.1 | 2.85 ± 0.02 |
| 0.52 | 2.93 ± 0.15 |

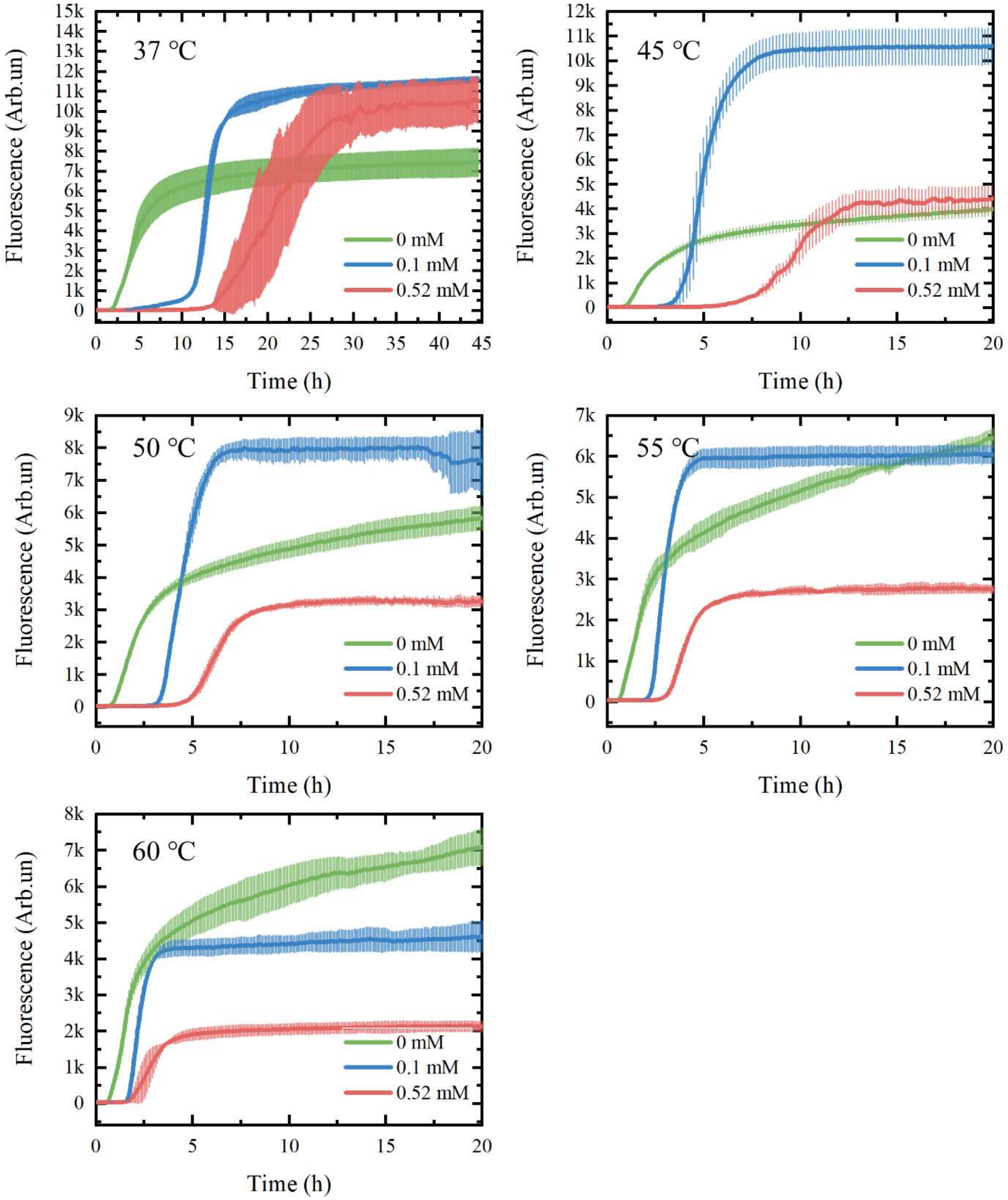


**Figure S1.** ThT fluorescence kinetics of insulin amyloid spherulites formation at different temperatures (mean±SD, n=4, except 0 mM at 50 °C, and 0.1 mM at 37 °C, where n=3). The concentration of added PS80 is shown as legend in the figures, while the temperature is shown in the top left corner of each panel.

**Differential Scanning Calorimetry (DSC)**

3 mg/mL insulin was dissolved in 20% (v/v) acetic acid with the addition of 0.5 M NaCl, with (0.52 mM) or without PS80 and filtered through a 0.22 µm cellulose acetate (CA). DSC scans were carried out using a VP-DSC differential scanning microcalorimeter with an autosampler (MicroCal, LLC, Northampton, MA, USA). 20% (v/v) acetic acid with the addition of 0.5 M NaCl with (0.52 mM) or without PS80 were used as reference. All scans were performed from 10 °C to 120 °C at a scan rate of 1 °C/min with nitrogen flow to keep the pressure constant.

**Spherulites Size Distribution at Different Temperatures**

3 mg/mL insulin was dissolved in 20% (v/v) acetic acid with the addition of 0.5 M NaCl, and with (0.52 mM) or without the addition of PS80. Filtered insulin solutions were incubated in microtubes at 32, 37, 45, and 60 °C for 240, 72, 48, and 48 hrs, respectively. For spherulites incubated at 45 °C, spherulites formed with 0.1 mM PS80 were also analyzed. After the aggregation reaction, samples were centrifuged for 10 min at 10 °C and 9279 g and the supernatant were separated from the pellets. The residual insulin in the supernatant was determined using a UV spectrophotometer (UV 1900, Shimadzu, Kyoto, Japan) using 1.0 $(mg/mL)^{-1}cm^{-1}$ as the extinction coefficient at 276 nm. The data are shown in Table S2.

**Table S2**. Residual insulin concentration (mg/mL) after the aggregation process at different concentrations of polysorbate 80 (PS80) and different temperatures (mean ± SD, n=2, N=2, n represents the different times of measurements, while N represents samples from different batches).

| Sample/Incubation Temperature | 32 °C | 37 °C | 45 °C | 60 °C |
|---|---|---|---|---|

| | | | | |
|---|---|---|---|---|
| 0 mM PS80 spherulite, mg/mL | 0.07 ± 0.05 | 0.02 ± 0.02 | 0.05 ± 0.04 | 0.06 ± 0.03 |
| 0.1 mM PS80 spherulite, mg/mL | 0.00 ± 0.01 | -0.03 ± 0.01 | -0.01 ± 0.01 | 0.01 ± 0.01 |
| 0.52 mM PS80 spherulite, mg/mL | 0.03 ± 0.00 | -0.03 ± 0.01 | -0.01 ± 0.01 | 0.04 ± 0.02 |

Microtubes containing the pellets (i.e. spherulites) were vortexed and the aggregates deposited on the microtube surface re-dispersed in solution. The samples were transferred by pipetting in and out 10 times before dilution in purified water (1:20 (v/v)) and addition of ThT (final concentration of ThT 20 µM). 40 µL of the diluted samples were transferred into the wells of a 96-well plate. For each sample, five replicates were prepared. The plate was gently tapped on the table to make sure the diluted samples volume covered the surface on the bottom of the wells. Measurements were performed 15 min after the plate preparation to allow for the precipitation of the aggregates on the well bottom. The plate was measured under matrix mode (5×5, Figure S2) with a Clariostar plate reader to obtain the ThT fluorescence distribution of each well. The two positions with the highest fluorescence values within each well were used for imaging and size distribution calculations (for all the samples, the middle of the well showed the highest fluorescence value). Images were taken using a Leica DMi8 optical microscopy (Wetzlar, Germany) using a 10× objective (Leica Microsystem, Wetzlar, Germany). The diameters of spherulites were measured via ImageJ[2]. Images of spherulites can be found in Figure S3.

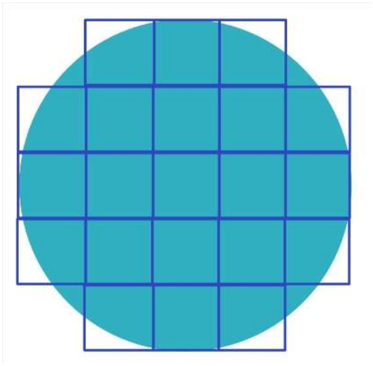

**Figure S2**: Scheme of a well that is measured under the 5×5 matrix mode. The squares represent the places where the fluorescence value is measured. Created with BioRender.com

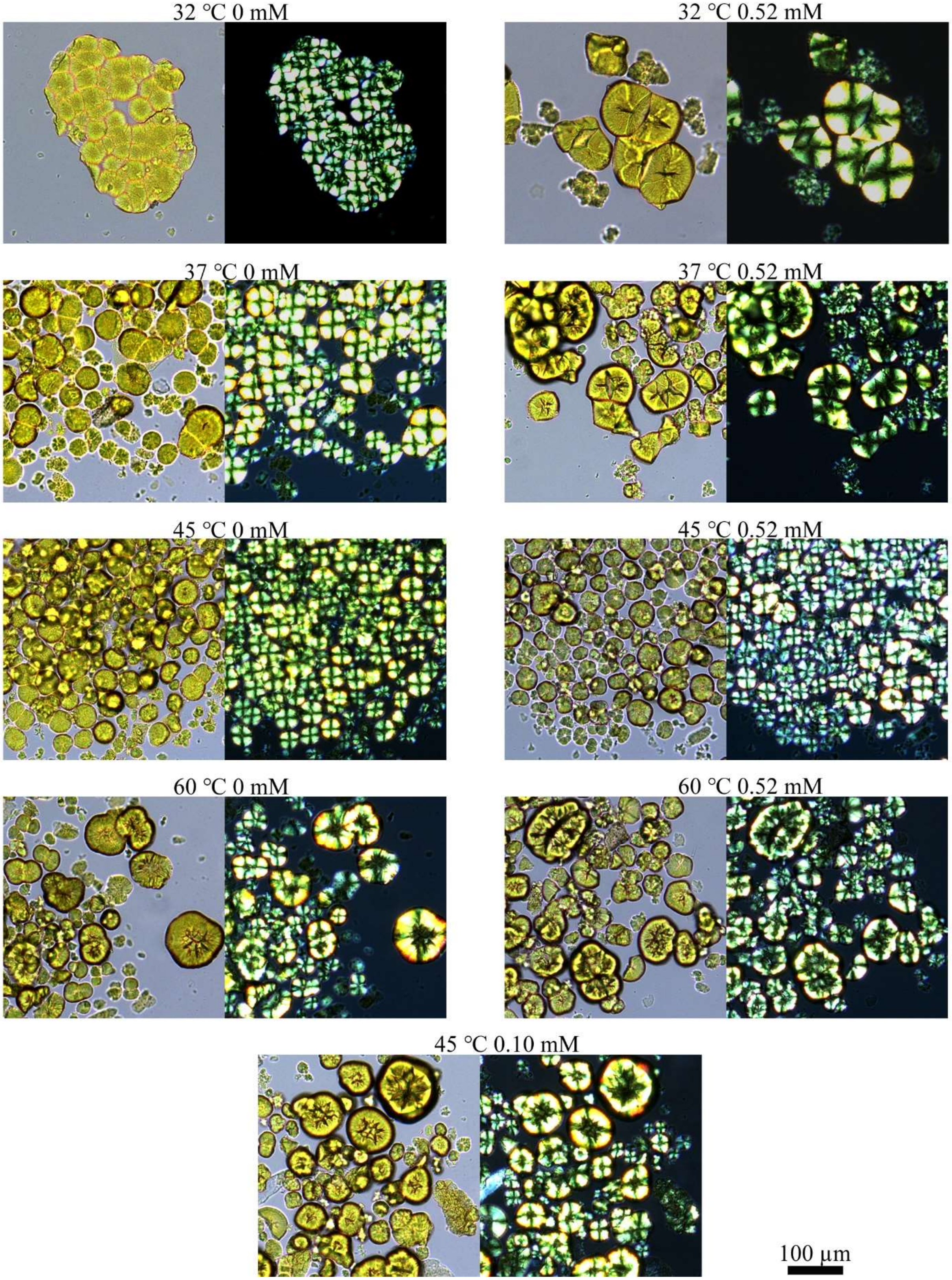
32 °C 0 mM
32 °C 0.52 mM
37 °C 0 mM
37 °C 0.52 mM
45 °C 0 mM
45 °C 0.52 mM
60 °C 0 mM
60 °C 0.52 mM
45 °C 0.10 mM
100 μm

**Figure S3.** Optical images of insulin spherulites incubated at different temperatures without and with different concentrations of polysorbate 80 (PS80). Images taken under bright field is shown on the left, while a Maltese cross can be detected under polarized light on the right, which is characteristic of spherulite formation. 3 mg/mL insulin was incubated in 20% (v/v) acetic acid and 0.5 M NaCl. Scale bar: 100 µm.

During the measurement, the longest distance/diameter through the spherulites centre was recorded (example, Figure S4a). Additionally, if the spherulites had a visibly blurry outline, the diameter was measured from the inner rims (example, Figure S4b). For the spherulites that formed clusters, only those which were distinguishable as a single entity (example, Figure S4c) and could be identified with the Maltese cross under polarized light were measured (example, Figure S5). The spherulites which were hidden in the clusters, or out of focus were not considered in the size distribution (example, Figure S5b).

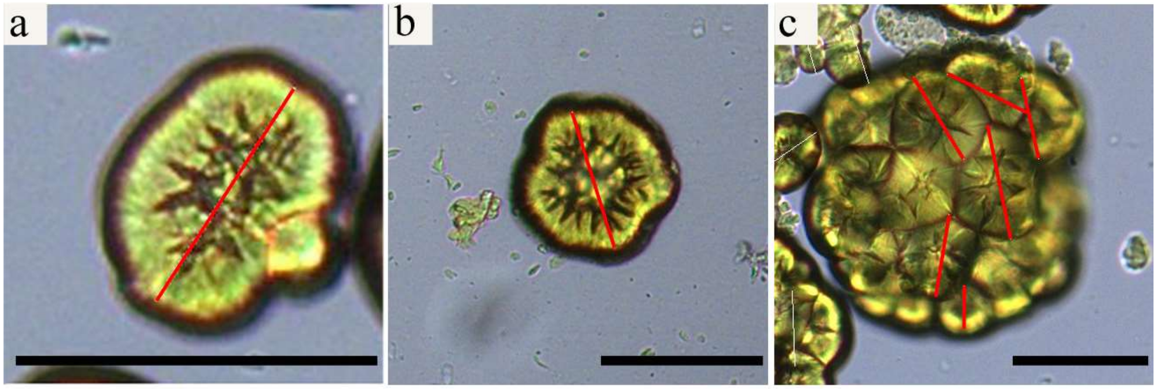


**Figure S4**. a) Example of an elongated spherulite and how its diameter was measured (red line). b) Example of a big spherulite and how its diameter was measured. c): Example of a spherulite cluster illustrating how only clearly distinguishable spherulites were selected to be included in the statistics. Scale bar: 100 µm.

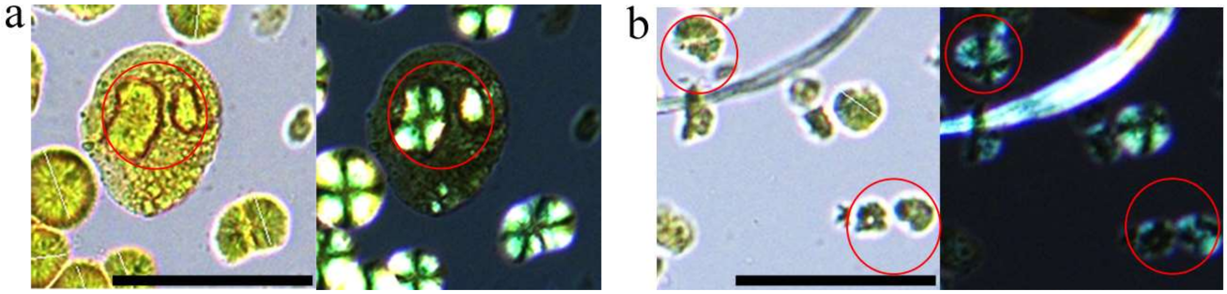


**Figure S5**: a) Example of a spherulite that is covered by other aggregates, but distinguishable by using the polarized light image, confirming the Maltese cross formation. b) Example of two spherulites, both showing Maltese cross formation, but lacking eligibility for measurement due to them being out of focus. Scale bar: 100 µm.

**Transmission Electron Microscopy (TEM)**

Copper grids (400 mesh, Agar Scientific, Stansted, UK) were first coated with Formvar and carbon film. 5 µL of the spherulites sample was loaded onto the grid and left for 60 s. 10 µL of distilled water was added on the grid, and excess water was removed using filter paper. 10 µL of 2% (v/v) uranyl acetate (Agar Scientific, Essex, UK) was added for staining for 30 s. Finally, 10 µL distilled water was used to wash 2 times, and, after each wash, the excess water was removed using filter paper. After drying, the grids were then ready to use. Images were collected on a Philips CM100 transmission electron microscope (Hillsboro, OR, USA), operating at an acceleration voltage of 80 kV.

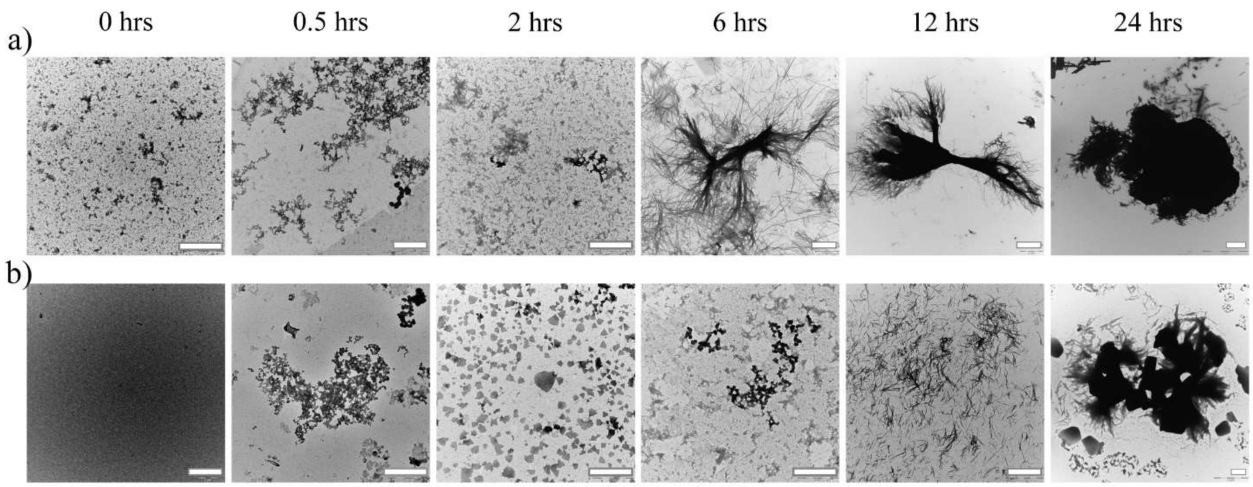


**Figure S6.** Transmission electron microscopy (TEM) images for insulin incubated in 20% (v/v) acetic acid in 0.5 M NaCl and with 0 µM (a) and 100 µM (b) polysorbate 80 addition. Samples were incubated at 45 °C and at the specified time the samples were taken out and analyzed via TEM. Scale bar: 2 µm.

TEM micrographs in Figure S6 confirm the aggregation delay induced by PS80 and the appearance of massive aggregation also observed via bulk kinetic analysis (Figure S1). Indeed, in the absence of PS80, early aggregates were observed in the first hour, potentially contributing to the formation of spherulites, which were observed after 6 hrs of incubation (Figure S6a). In agreement with a recent report from our group[3], the formation of early bundle-like fibrils was observed after 6-12 hrs of incubation (Figure S6a). This type of morphology is indicated as intermediate species eventually maturing into spherical spherulites. In the presence of PS80, spherulites only appear at ~ 24 hrs (Figure S6b) and only dispersed fiber-like materials were detected in the early phases of the process by TEM. Interestingly, spherulites were not the sole species along the pathway and even upon the completion of both processes, the presence of filamentous fiber-like material is evident. This confirms the possibility of the co-existence of different species within the same protein ensemble[4-7].

### ThT Kinetics Recorded in the 5×5 Grid Setup

Similar to the matrix grid setting mentioned above (please see section *"Spherulites Size Distribution at Different Temperatures"*), the ThT fluorescence value of a sample was set to be measured in the 5×5 grids per well. The kinetics were obtained by conducting the fluorescence 5×5 matrix scan per well every 15 min at 45 °C in the Clariostar plate reader. The setting allowed us to obtain 21 kinetic traces per well. Samples of 3.0 mg/mL insulin in 0, 0.1, and 0.52 mM PS80 solutions were prepared as previously described (please see section "*Thioflavin T Fluorescence (ThT) Kinetics"*). The addition of ThT and preparation of the samples in the 96 well plate was done as previously described. 4 wells of each sample were included in the data analysis process. The 0 mM PS80 sample measurement was performed immediately after the solutions were placed in the plate reader system and lasted for 7.5 hrs (the plateau was reached at around 5 hrs). 0.1 mM PS80 and 0.52 mM PS80 samples were firstly incubated in the plate reader at 45 °C for approximately 8.5 hrs (no data acquisition in the lag time). After 8.5 hrs, data collection was initiated and lasted for approximately 12 hrs. Once the plateau (saturation phase) was reached, measurements were stopped, but the plates were allowed to incubate throughout the night. The next day, a final ThT fluorescence 5×5 grid scan was measured. In total, the 0 mM PS80 insulin sample samples were incubated for approximately 28 hrs, while the 0.1 mM and 0.52 mM PS80 samples were incubated for approximately 59 hrs. This setup was necessary since the measurements were retrieved manually.

### Real-time Kinetic via Photobleaching Localization Microscopy (REPLOM)

Insulin used in this experiment was labelled by Alexa Fluor 647 NHS ester (ThermoFisher Scientific). 5 μL of 2 mg/mL Alexa Fluor 647 NHS ester dissolved in anhydrous-DMSO was added into 1 mL of a 5 mg/mL insulin solution (in PBS, pH 7.4), the solution was equilibrated at room temperature for around 2 hrs for conjugation. The labelled insulin was then purified via a PD SpinTrap G-25 column (GE Healthcare, Chicago, IL, USA) and diluted to a final concentration of 1 mg/mL. The aliquots were stored at -80 °C.

3 mg/mL insulin in 20% (v/v) acetic acid in 0.5 M NaCl, with (0.1 mM) or without PS80 incubated in microtubes on a heating block at 45 °C for around 7 hrs. Before the incubation, 3 μL of the 10 times diluted labelled insulin was added into dissolved insulin solution and the whole solution filtered via a 0.22 μm CA filter. The molar ratio between labelled and unlabelled insulin was 1 to 10,000.

After 7 hrs, the solutions were transferred to the microscope chamber on a poly-L-lysine coated glass surface. Microscope cover slides (Hecht Assistent®, Sondheim vor der Rhön, Germany) after cleaning (2% (v/v) Hellmanex®, purified water and methanol, sequentially, and with 3 times of 10 min sonication at each step) were stored in methanol. A plasma cleaner was used to further clean the cover slides for 2 min before affixing to the flow chambers (sticky-Slide VI 0.4, ibidi GmbH, Munich, Germany). 0.01% (w/v) poly-L-Lysine (Sigma-Aldrich, Burlington, MA, USA) was added. After 2 hrs of incubation, the chambers were washed 3 times with purified water and dried with nitrogen.

REPLOM was conducted on an inverted optical microscope (Olympus IX-83, Tokyo, Japan) by using a 100× oil immersion objective (UAPON 100XOTIRF, NA=1.49, Olympus) and cellVivo incubation system (Olympus, Tokyo, Japan). The excitation of Alexa Fluor 647 was performed at 640 nm with a solid state laser line (Olympus). The signal of Alexa Fluor 647 was recorded with an EMCCD camera (imagEM X2, Hamamatsu Photonics K.K., Shizuoka, Japan) with an exposure time of 30 ms. The waiting time for each image was 27.3 s for spherulites samples without PS80, and 30 s for samples containing 0.1 mM PS80. The incubation temperature was 45 °C maintained by a heating unit 2000 (PeCon GmbH, Erbach, Germany).

The spherulite growth was identified by the fluorophores bound on the insulin molecules, images and growth rate fitting were obtained according to the method developed in our previous work as well as the criteria used for including the data in the overall statistics[3].

Video recordings of spherulites growth are found in Video S1 (isotropic growth) and Video S2 (anisotropic growth) as examples.
